# ADAM10- and presenilin 1/γ-secretase-dependent cleavage of PTPRT mitigates neurodegeneration of Alzheimer’s disease

**DOI:** 10.1101/2021.01.11.426157

**Authors:** Siling Liu, Zhongyu Zhang, Lianwei Li, Li Yao, Zhanshan Ma, Jiali Li

## Abstract

PTPRT (receptor-type tyrosine-protein phosphatase T), as a brain-specific type 1 transmembrane protein, plays an important function in neurodevelopment and synapse formation. However, whether PTPRT-dependent signaling is involved in Alzheimer’s disease (AD) remains elusive. Here, we identified that PTPRT intracellular domain (PICD), which was released from ADAM10- and presenilin 1-/γ-secretase-dependent cleavage of PTPRT, efficiently translocated to the nucleus via a conserved nuclear localization signal. Inhibition of nuclear localization of PICD via the mutation of its nuclear localization signal (NLS) leads to accumulation of phosphorylated signal transducer and activator of transcription 3 (pSTAT3), which is a substrate of PTPRT and eventually resulted in neuronal cell death. Consistently, RNA sequencing reveals that expression of the PICD alone can profoundly alter the expression of genes associated with synapse function and dephosphorylation, phosphatase and cell adhesion. Unexpectedly, the downregulated levels of *Ptprt* mRNA and protein were found in both human and mouse AD brains. Finally, overexpression of PICD alone not only significantly decreases the level of phosph-STAT3^Y705^ and Aβ deposition in the hippocampus of APP/PS1 mice, but also improves synaptic function and behavioral deficits in APP/PS1 mice. Our findings suggest that a novel role of the ADAM 10- and presenilin 1-/γ-secretase-dependent cleavage of PTPRT in the events can mitigate neurodegeneration of AD and moderate Alzheimer’s pathogenesis.

## Introduction

Extracellular deposition of the amyloid-β peptide is a major hallmark pathology for both early-onset and late-onset Alzheimer’s disease, and it has been taken as the culprit in the appearance of synaptic dysfunction, neuronal degeneration and loss according to the amyloid hypothesis. The amyloid-β peptide (Aβ) is a fragment of a large type I membrane protein known as the amyloid precursor protein (APP). Mechanically, Aβ is released from APP by the combined actions of shedding by β secretase (BACE1) in the juxtamembrane region and the followed cleavage by the γ secretase/presenilin-1 complex in the intramembrane domain, resulting in secretion of the neurotoxic amyloid-β peptide and a cytoplasmic peptide known as the APP intracellular domain (AICD)[1]. In addition to APP, some other type 1 transmembrane proteins are subject to similar processing, which is also called regulated intramembrane proteolysis (RIP)[2]. RIP, as an evolutionally conserved signaling mechanism, is very important for organisms[2]. The p75 receptor, another important γ-secretase substrate, is also affected in AD pathogenesis as the released intracellular domain can induce neuronal apoptosis after translocating to the nucleus[3, 4]. TREM2, another type 1 transmembrane protein and AD risk-factor gene, can also be cleaved by α/γ-secretase action[5]. The ectodomain sTREM2 released by ADAM10 induces inflammatory responses and enhances microglial survival[6, 7], while the function of the intracellular domain is uncertain. RIP mediated signaling is also important for neurodevelopment, as illustrated by yet another γ-secretase substrate, Notch. After binding with its ligands, cleavage of Notch liberates its intracellular domain (NICD), which enters the nucleus to regulate developmental gene expression. The Notch pathway has been shown to regulate neurogenesis, synaptic plasticity and long-term memory[8–10]. Sequential cleavage of EphA3 by α- and γ-secretases mediates axon elongation, and the released intracellular domain of EphA3 is essential for the process[11].

PTPRT, a receptor-type protein tyrosine phosphatase, is a type 1 transmembrane protein with two intracellular catalytic domains. It is predominantly expressed in the brain of mammals. Clinical and sequencing studies have shown that mutations in the *Ptprt* gene would cause intellectual development and neurological disorders[12]. Several GWAS studies show that *PTPRT* is heavily involved in neurodevelopmental disorders[12–15]. PTPRT works as a regulator for synaptic formation via its dephosphorylation activity[16]. PTPRT could also regulate dendritic arborization and fusion of synaptic vesicles by dephosphorylating BCR and Syntaxin-binding protein 1 during neurodevelopment[17]. At the molecular level, PTPRT plays a crucial role in synapse formation by interacting with cell adhesion molecules[16]. Dephosphorylation of BCR protein by PTPRT regulates neuronal dendritic arborization[17].

PTPRT is also a potent phosphatase for pSTAT3 at Y705 residue[18]. STAT3, as a transcriptional factor, responds promptly to environmental stimulation by internal and external changes, which is important for cellular adaption[19, 20]. The rapid coordination between tyrosine kinases and phosphatases is important for the process. After being phosphorylated at Y705 by JAK, pSTAT3^Y705^ forms dimers and then accumulates in the nucleus, activating the transcription of a wide range of genes, including some oncogenes. Mutations or epigenetic silencing in the *Ptprt* gene could promote tumor growth due to the hyperactivation of STAT3[21, 22]. In the central nervous system, STAT3 also plays essential roles. STAT3 modulates neurogenesis and differentiation during brain development[23, 24], and activation of STAT3 is neuroprotective against brain ischemia by upregulating survivin and Mn-SOD[25]. In neurodegenerative diseases, neural STAT3 activation is a common phenomenon, and more evidence has showed its detrimental role for neuronal survival[26]. Activated STAT3 has also been found to facilitate long-term depression (LTD) in the AD mouse model and promote β-amyloid production by upregulating *BACE 1* expression in neurons[27].

In this study, we found that the downregulated mRNA and decreased protein levels of PTPRT led to the accumulation of pSTAT3^Y705^ in the brains of human AD and model mice. We further identify that PTPRT, a regulator of STAT3 in the mammalian brain, is a novel substrate for ADMA10 and presenilin 1/γ-secretase. After cleavage by ADAM10 and presenilin 1/γ-secretase, the intracellular domain of the released PTPRT (PICD) is translocated to the nucleus and regulates gene expression via dephosphorylation of pSTAT3^Y705^. Overexpression of PICD alone significantly reduces pSTAT3^Y705^ and Aβ deposition, and improves synaptic function and behavioral deficits in APP / PS1 mice.

## Materials and Methods

### Antibodies and Reagents

Antibodies against PTPRT (MABS1158 for WB and HPA017336 for IHC) and Flag (F3165) were from Sigma; pSTAT3 (Y705) (AP0070), ADAM10 (A10438) and ADAM17 (A0821) antibodies were from ABclonal; antibodies against Aβ_42_ (ab201061), STAT3 (ab68153), α-tubulin (ab7291), β-actin (ab8227), Presenlin-1 (ab15456), VAMP2 (ab3347) and Map2 (ab32454 and ab11267) were obtained from Abcam; antibodies against cleaved-Caspase3 (#9661), a-tubulin: (#3873S), histon3 (#9715) were purchased from Cell Signaling Tech; antibody against the Aβ plaque 6E10 (SIG-393200) antibody were from Covance; antibody against oligomeric Aβ MOAB-2 (NBP2-13075) and biotin-labeled MOAB-2 (NBP2-13075B) from Novus Biologicals. The secondary antibodies used for immunocytochemistry were as follows: chicken anti-mouse or rabbit Alexa 488; donkey anti-mouse or rabbit Alexa 594 (Invitrogen, Eugene, OR); all were used at a dilution of 1:500. ProLong™ Diamond Antifade Mountant with DAPI (Thermo Fisher Scientific, P36962) was used as a nuclear counterstain at 1 µg/ml. DAPT (γ-secretase inhibitor, D5942), Thioflavine S (T1892), Anti-Digoxigenin-AP (Fab fragments, 11093274910), and Phorbol 12- myristate 13-acetate (PMA, p8139) were ordered from Sigma-Aldrich.

### Constructs

PLV-Ptprt-FL-mCherry and PLV-Ptprt-ICD-mCherry were constructed from the backbone of pLV-mCherry (Addgene, #36084). AAV-hSyn1-nls-luciferase-FLAG (control) and AAV-hSyn1-Ptprt-ICD-FLAG were constructed from the backbone of AAV-hysn1-GCaMP6s-P2A-nls-dTomoto (Addgene gene, # 51084). AAV-hysn1-GCaMP6s-P2A-nls-dTomoto was digested with BamH1 and EconR1. nls-luciferase-FLAG and PTPRT-ICD-FLAG were PCR amplified separately and the ligated with the backbone after BamH1 and EconR1 digestion to make AAV-nls-luciferase-FLAG and AAV-hSyn1-Ptprt-ICD-FLAG. nls and FLAG sequence were added by PCR amplification. pcDNA3.1-Ptprt was cloned from mouse brain RNA by SuperScript III one-step RT–PCR system (Invitrogen). Lentiviral shRNAs against mouse ptprt were constructed in pLKO.1 (Invitrogen) and transfected with pMD2g and psPAX into HEK293T cells for viral package. The sequences of shRNA targets can be found in Supplementary Table 1.

### Viral Preparation

Lentivirus preparation, the construct was co-transfected to HEK293T cells pMD2g and psPAX. The original medium was changed with the fresh one. 48 hours after that, the medium was collected. The lentivirus particles were concentrated by centrifuging at 25,000 rpm for 2 hours. The production of lentiviral particles up to 109 IU/ml was purified by Ultra-Pure Lentivirus purification kits (Applied Biological Materials Inc. Vancouver). AAV preparation, the constructs were packaged into AAV9 by the MIRCen viral production platform. Briefly, viral particles were produced by transient co-transfection of HEK-293T cells with an adenovirus helper plasmid, an AAV packaging plasmid carrying the rep2 and cap9 genes, and the AAV2 transfer vector containing the above-mentioned expression cassettes by PEI. 72 h following transfection, virions were purified and concentrated from cell lysate and supernatant by ultracentrifugation on an iodixanol density gradient followed by buffer exchange to PBS, 0.01 % Pluronic via a 100kd Amicon Centrifugal filter unit (Merck-Millipore, Darmstadt, Germany). Concentration of the vector stocks was estimated by quantitative PCR and expressed as viral genomes per ml of concentrated stocks (vg/ml).

### Human brain tissues

Human autopsy paraffin-embedded 10 μm brain sections were from the following sources with approval from the appropriate local regulatory authorities. We examined 16 case-patients graciously provided by the University of Pittsburgh Alzheimer’s Disease Research Center (ADRC) brain bank with approval from the Committee for Oversight of Research and Clinical Training Involving Decedents (CORID). Basic information has been described as our previous study[28]. Additional frozen tissue was a generous gift of the ADRC at Washington University in St. Louis (Grant P50-AG-05681) with approval from the Neuropathology Core (protocol #T1016) at Hongkong University of Science and Technology.

### Animals

Animal studies were performed with age- and sex-matched wild-type and APP/PS1 mice. APP/PS1 mice carry two different human genes harboring disease-associated mutations: PS_1m146v_/APP_swe_. Breeding stock was purchased from The Jackson Laboratory. Animals were housed and bred at the animal facility of the Kunming Institute of Zoology. All animal experiments were conducted under license from the Kunming Institute of Zoology and were approved by the Animal Care and Use Committee of the Kunming Institute of Zoology, Chinese Academy of Sciences (SMKX-SQ-20170628).

### Bioinformatic analysis

For human PTPRT expression analysis, we followed the method from a published paper[29]. Briefly, 20 GSE series of expression data from 684 AD and 562 control human postmortem brain samples was re-normalized, and then the expression of PTPRT was extracted and analyzed. For correlation analysis between AD pathological burden and the level of PTPRT expression, a GEO dataset GSE64398 was used[30], and simple linear regression analysis was conducted. To analyze the PTPRT expression in the hippocampus of AD patients at various stages of severity, we used GEO dataset GSE1297[31].

### Mouse embryonic fibroblasts (MEFs), N2a, and HEK293T cell culture

All the cells (MEF, N2a and HEK293) were plated onto tissue culture dishes in DMEM (Invitrogen) supplemented with 10% FBS and 1% Pen/Strep (Gibco) at 37°C and 5% CO2. PEI (1 mg/ml) was used for the transfection of HEK293T (WT or mutant) cells. Lentivirus infection of MEF and N2a cells was performed at the second day of subculture when the cell confluence reached 70%. Polybrene (final concentration is 8 µg/mL) was used to facilitate the infection.

### RNA extraction and reverse transcription PCR

Total RNAs from N2a cells transfected with either pLV-mCherry or pLV-Ptprt-ICD-mCherry were extracted and prepared using PureLink micro-to-midi total RNA purification system (Invitrogen). Quantitative Real-time PCR, cDNA was prepared by using oligonucleotide (dT), random primers, and a Thermo Reverse Transcription kit (Signal way Biotechnology). The mRNA level of *Gapdh* was used as an internal control. The sequences of qPCR primers can be found in Supplementary Table 2.

### Primary Neuronal Cultures

Mouse embryonic cortical neurons were isolated by standard procedures. All embryonic pups from APP/PS1 x WT mating were harvested and treated separately then retrospectively genotyped by PCR. Isolated E16.5 embryonic hippocampus was treated with 0.25% Trypsin-EDTA and dissociated into single cells by gentle trituration. Cells were suspended in Neurobasal medium supplemented with B27 and 2 mM glutamine, then plated on coverslips or dishes coated with poly-L-Lysine (5 mg/mL). The cells were seeded at 1×10^4^ cells/cm^2^. All cultures were grown for a minimum of 7 days *in vitro* (DIV) before any treatment. All viral infections were carried at DIV5.

Primary neurons were infected using a multiplicity of infection (moi) between 5 and 10 to provide an efficiency of infection above 70%. All data were collected and analyzed in a double-blind manner.

### Immunohistochemistry

For DAB/bright field staining, all paraffin-embedded human sections were deparaffinized in xylene and then rehydrated through graded ethanol to water. The sections were pretreated with 0.3% hydrogen peroxide in methanol for 30 min to remove endogenous peroxidase activity, rinsed in Tris-buffered saline (TBS), and then treated with 0.1 M citrate buffer in a microwave at sufficient power to keep the solution at 100°C for 20 min. Sections were cooled in the same buffer at room temperature (RT) for 30 min and rinsed in TBS. Slides were incubated in 10% goat serum in PBS blocking solution for1 h at RT, after which the primary antibody was applied to the sections that were then incubated at 4°C overnight. The sections were washed three times in TBS before applying the secondary antibody for 1 hour at RT. Afterward, sections were rinsed and incubated in Vectastain ABC Elite reagent, and developed using diaminobenzidine according to the protocol of the manufacturer (Vector Laboratories). The sections were counterstained with hematoxylin and mounted in Permount. Control sections were subjected to the identical staining procedure except for the omission of the primary antibody. All sections were analyzed and imaged by using AxioImager. Z1 (Zeiss), and LSM 780 NLO confocal (Zeiss), microscopes. The collection and analysis of all data were performed in a double-blind manner.

### Immunofluorescence

All paraffin-embedded human sections were deparaffinized in xylene and then rehydrated through graded ethanol to water and PBS. For the mouse, cryostat sections were first rinsed in PBS, Subsequent steps were identical for paraffin and cryostat material. All primary antibodies were diluted in PBS containing 0.5% TritonX-100 and 5% goat serum and incubated with sections overnight at 4°C. After rinsing in PBS, they were incubated for 1 hour with the appropriate secondary antibody, which was conjugated with a fluorescent label. All sections were mounted in ProLong® Gold anti-fade reagent with DAPI (Invitrogen) under a glass coverslip. All immunofluorescence images were collected using a Zeiss Olympus IX-81 microscope with either a 20× or 40× objective running Metamorph. Microsoft Excel was used to calculate the fraction of positive clusters. GraphPad Prism was used to perform statistical analysis and to visualize bar charts. Error bars represent S.E.M. All data were collected and analyzed in a double-blind manner.

### Protein extraction and western blot

Brain tissues were lysed using 1×lRIPA buffer (Sigma Aldrich), complemented with PIC (Protease Inhibitor Cocktail, Cat #11836153001, Roche) and Phosphatase Inhibitor Cocktail (Cat #11836153001, Roche). After protein quantification, samples were boiled for 10lmin in SDS loading buffer (1:1 ratio). An aliquot (up to 50lμg) of the resulting sample was run on an SDS–PAGE gel and then transferred to a Hybond-PVDF membrane (Amersham Pharmacia). The membrane was blocked in 5% milk for 1 hour at room temperature and incubated overnight with the appropriate primary antibody re-suspended in 5% milk in TBST at 4l°C. Following three washes in TBST, the membrane was incubated for 2lhours at room temperature with the appropriate secondary antibody re-suspended in 5% milk in TBST at room temperature, followed by 4 washes using TBST. Signal ECL (Pierce) amplification was detected by Tanon 5200 Multi chemiluminescent image system (Tanon, Shanghai, China). Signal intensity was quantified with Image J (National Institute of Health).

### RNA sequencing

Libraries were prepared from the RNAs of three 3 biologically independent experiments. Sequence-libraries of each sample were equimolarly pooled and sequenced on an Illumina NextSeq 500 instrument (High Output, 75 bp, Single Reads, v2) at the core facility of the Kunming Institute of Zoology. Differentially expressed genes were uploaded into the IPA software (Ingenuity Systems, http://www.ingenuity.com). An IPA core analysis was performed focusing on both up- and down-regulated molecules and setting the log ratio (LR) cutoff ≥ 0.58 (fold change ≥ 1.5) and the FDR cutoff ≤ 0.0005 (range 0.0–0.0005). FastQC (v0.119) and Fastx-Toolkit (v0.0.13) were used to clean the sequencing data. We use Tophat (v2.1.1) to align clean sequencing reads to the exon of the reference genome (Mus musculus (house mouse), GRCm38.p6). The unmapped reads were split into shorter fragments and then aligned to splice junctions between exons. Cufflinks (v2.1.1) use the alignment files from Tophat to assemble and reconstruct the transcriptome. The transcript abundances were estimates by Cuffnorm and differential expression of transcripts performed by Cuffdiff.

### Data availability

RNA-seq raw data have been deposited in NCBI’s Gene Expression Omnibus and are accessible through GEO Series accession number GSE159943 (https://www.ncbi.nlm.nih.gov/geo/query/acc.cgi?acc=GSE159943).

### Virus production and stereotactic intracranial injection

Viral preparation and hippocampal microinjection were conducted as previously described[28]. Briefly, the high adenoviral PICD viral particle titers were prepared with the Adenoviral Expression system and purified by Adenoviral Purification Kits (Applied Biological Material Inc. Vancouver). Stereotaxic intra-hippocampal infusions were delivered to wild-type and APP/PS1 mice (9 months of age) under isoflurane anesthesia. Mice were positioned in a Kopf stereotaxic apparatus and burr holes drilled into the skull. A 5-ml Hamilton syringe fitted with a 33-gauge needle was lowered into the hippocampus for viral particles delivery. The coordinates relative to bregma: anteroposterior, -2.18 mm; mediolateral, ±2.30 mm; and dorsoventral, -2.10 mm. Each hippocampus was injected with 1 μl of the adenovirus-associated viral particles using a Hamilton syringe. The injection rate was 0.2 μl/min. All experiment as indicated above for the mice were carried out 3 months after injection. All data were collected and analyzed in a double-blind manner.

### ELISA

Aβ extraction was performed on either micro-dissected brain tissue enriched for hippocampus from sex-matched 12-month-old wild type and APP/PS1 mice or DIV14 primary hippocampal neurons. Lysates were mixed with an equal volume of 0.4% diethylamine in NaCl and centrifuged for 14,000 *g* for 1 h at 4°C. The supernatant was collected and neutralized with 0.5 M Tris, pH 6.8, and analyzed as the soluble Aβ fraction. The pellet was sonicated with 70% formic acid and centrifuged at 100,000 *g* for 45 min at 4°C, and the supernatant was neutralized and analyzed as the insoluble Aβ fraction. Samples were analyzed by sandwich ELISA using mouse monoclonal MOAB-2 anti-Aß capture antibody, biotinylated MOAB-2 and Aß_42_ detection antibodies. All data were collected and analyzed in a double-blind manner.

### Field potential recording and LTP

Preparation and recording of hippocampal brain slices were described as previously[32]. Briefly, mice of 12-month-old wild type and APP/PS1 one week after completion of behavioral testing are decapitated under isoflurane anesthesia, and the brains removed. Extracellular recordings of field excitatory postsynaptic potentials (fEPSPs) were made with artificial cerebrospinal fluid (ACSF)-filled glass electrodes (5–10 mm tip diameter). Test stimuli (0.1 ms) were delivered with a bipolar platinum/iridium stimulating electrode at 1 min intervals, except for specialized protocols that elicited changes in synaptic strength (see below). For recordings of CA1 activation by Schaffer collateral stimulation, recording and stimulating electrodes were both placed in stratum radiatum. Each experiment was begun by obtaining input-output relationships to establish the strength of baseline synaptic transmission. A Grass S8800 stimulator connected to a Grass PSIU6 photoelectric stimulus isolation unit was used to deliver a series of increasing-intensity constant-current pulses. Current magnitude was adjusted to elicit responses ranging from just suprathreshold to near maximal. Following this, stimulus intensity was adjusted to evoke fEPSPs 30%–40% of maximum, typically 30–40 mA. To elicit LTP, TBS was used. A single TBS consisted of 12 bursts of four 100 Hz pulses spaced 200 ms apart. Response magnitude was quantified with the slope of the field potential. Statistics Test of significance were performed using either ANOVA, or paired and unpaired *t*-tests as appropriate.

### Morris Water Maze

WT mice and APP/PS1 mice in the AAV-control and PICD groups were trained in an open circular pool (diameter: 170 cm) filled with water at a depth of 30 cm and maintained at 22 °C ± 1°C as described previously[33]. For data collection, the maze was divided into four equal quadrants (NE, SE, SW and NW) by designating two orthogonal axes, the end of which demarcated four cardinal points: north, south, east, and west. Mice that failed to find the platform within 60 s were placed on it for 10 s, the same period as was allowed for the successful animals. White geometric figures, one hung on each wall of the room, were used as external visual clues. Behavior was evaluated by direct observation and analysis of videotape-recorded images. Variables of time (escape latency) and quadrant preference and entries were analyzed in all the trials of the tasks. The escape latency was readily measured with a stopwatch by an observer unaware of the animal’s genotype and confirmed during the subsequent video-tracking analysis. In the probe trial, the time spent and number of entries in each of the four quadrants were also measured retrospectively by means of the automated video-tracking analysis.

### Statistics

All data were obtained under the same conditions. And all experiments and analysis were performed in a double-blind manner. No randomization was performed to allocate subjects in the study. No sample calculation was performed. No exclusion criteria were pre-determined. No animal was excluded based on the exclusion criteria or died during experiments. Data were presented as the mean ± S.E.M. *P* values describing significance were based on Student *t-*tests (2-tailed; α = 0.05) or analysis of repeated-measures two-way ANOVA. *P* values less than 0.05 indicate statistical significance. GraphPad Prism 8.0 was used to perform ANOVAs and *t*-tests and to visualize bar charts.

## Results

### PTPRT is a novel substrate of ADAM10- and presenilin 1/γ-secretase

AS a type 1 transmembrane protein, PTPRT is a potential substrate of *γ* secretase. while the extracellular domain shedding in the juxtamembrane region is necessary for the subsequent cleavage by presenilin 1/γ-secretase complex. To determine whether PTPRT could undergo sequential processing, we treated the HEK293 cells overexpressed with C terminal FLAG-tagged PTPRT (intracellular domain) with different compounds. As is shown in the results, DAPT (γ-secretase inhibitor) could increase the accumulation of the smaller C terminal region fragment (CTF2) while it didn’t affect the bigger size C terminal region fragment (CTF1), which is already known to be produced by furin cleavage. PMA, an α-secretase activator via the PKC pathway, could increase CTF2 accumulation after DAPT inhibition. The treatment with ADAM/matrix metalloproteinase inhibitor GM6001 could decrease the CTF2 accumulation (Fig. 1a-b). Treatment with proteasome inhibitor MG132 solely helped us detect the accumulation of intracellular fragments (PICD), which is a little bit smaller than CTF2 (Red arrow represents PICD). We further examined endogenous PTPRT cleavage in primary neurons with an antibody against the PTPRT C-terminal. As expected, a similar result was found (Fig. 1c-d). ADAM10 and ADAM17 are the two most important α-secretases responsible for the ectodomain shedding. We tested γ-secretase cleavage in ADAM10 KO and ADAM17 KO HEK293 cells[34], respectively. As we could see, CTF2 didn’t accumulate significantly after DAPT treatment in ADAM10-KO cells compared with WT cells (Fig. 1e-f). It suggested that ADAM10 rather than ADAM17 is the major secretase for PTPRT shedding. For further verification of PTPRT cleavage by γ-secretase, we transfected wild type (MEF-WT) or *Psen1*^-/-^/*Psen2*^-/-^ double knockout mouse embryonic fibroblasts (MEF-DKO) with mouse *Ptprt* construct. PTPRT CTF2 was absent in WT cells while present in DKO cells (Fig. 1g-h). As the main component of the γ-secretase complex for intramembrane proteolysis, presenilin 1 is found to cut γ-secretase substrates without the help of presenilin 2. We expressed *Psen1* delivered by lentivirus to see if it could recuse the processing. Indeed, PTPRT CTF2 could not be detected in *Psen1* expressed MEF cells. Furthermore, γ-secretase could cleave PTPRT and generated an intracellular domain (PICD) at 37℃ in plasma membrane separated from *Ptprt-*overexpressed HEK293 cells; and the processing could also be inhibited by DAPT in which the PICD signals were diminished in contrast to an increased level of CTF2 (Figure S2a-b). Together, those results demonstrate the sequential cleavage of PRPRT by ADAM10 and presenilin1/γ-secretase.

**Fig 1.**
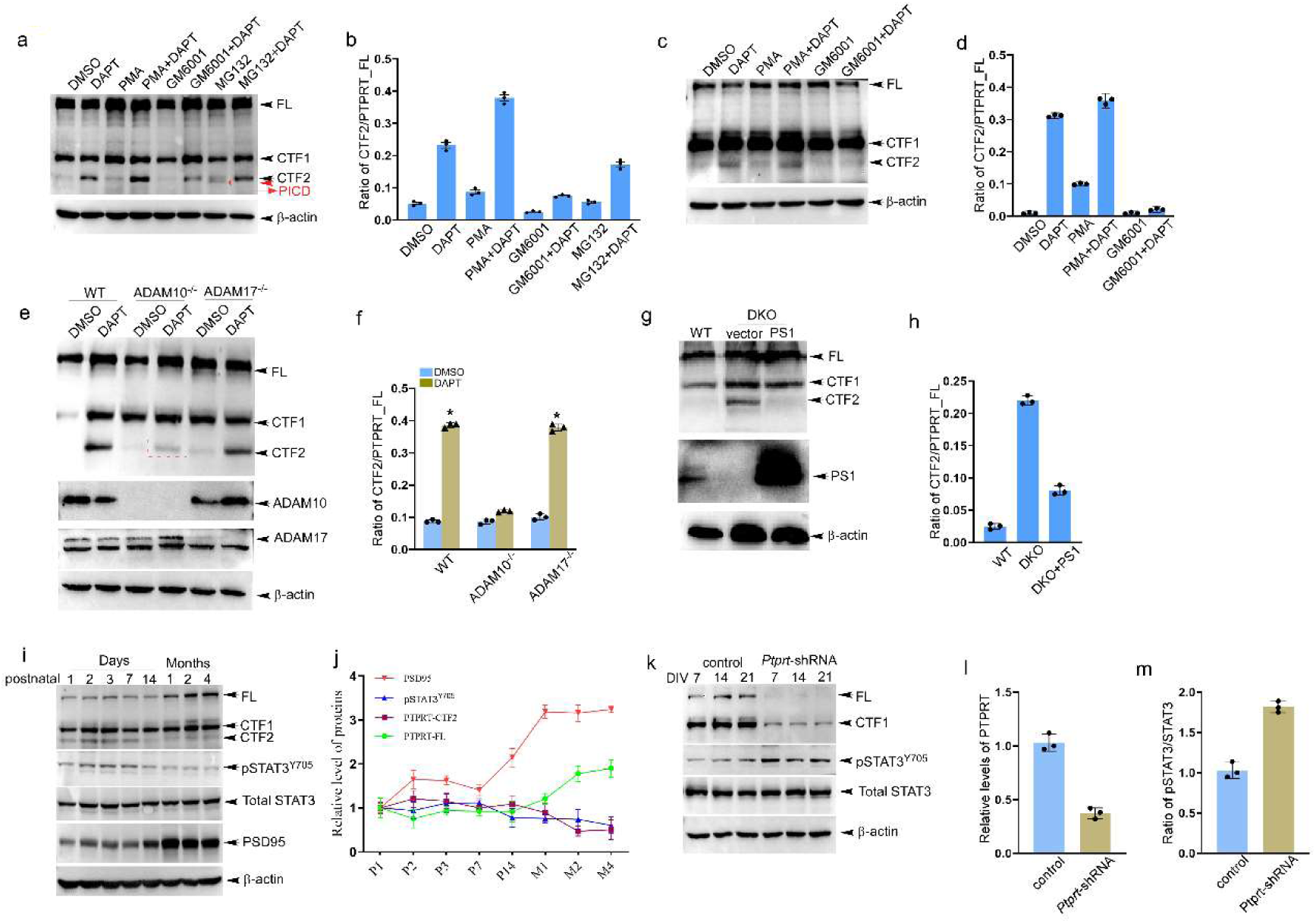
PTPRT was subsequently cleaved by ADAM10 and presenilin 1/γ-secretase. **a.** Protein extracts from HEK 293 cells with indicated pretreatments were assayed by Western blot for the presence of PTPRT and its fragments CTF1, CTF2. **b.** Quantification of the immunoblot intensities of PTPRT shown in panel a). Error bars denote S.E.M. (n= 3 independent experiments; *, *P* < 0.001). **c.** Protein extracts from primary cortical neurons of the wild type at DIV 14 with indicated treatment were assayed by Western blot for the presence of PTPRT and its fragments. **d.** Quantification of the immunoblot intensities of PTPRT shown in panel c). Error bars denote S.E.M. (n= 3 independent experiments; *, *P* < 0.001). **e.** Protein extracts from wild type and ADAM10^-/-^ and ADAM17^-/-^ HEK293 cells w/o DAPT treatment were assayed by Western blot for the presence of PTPRT and its fragments. **f.** Quantification of the immunoblot intensities of PTPRT shown in panel e). Error bars denote S.E.M. (n= 3 independent experiments; *, *P* < 0.001). **g.** Protein extracts from presenilin 1 and 2 DKO MEFs with overexpressing PS1 (presenilin 1) were assayed by Western blot for the presence of PTPRT and its fragments. **h.** Quantification of the immunoblot intensities of PTPRT shown in panel g). Error bars denote S.E.M. (n= 3 independent experiments; *, *P* < 0.001). **i.** Protein extracts fresh hippocampal tissues of wild type mice at indicated ages were assayed by Western blot for the presence of PTPRT and its fragments, and pSTAT3^Y705^. **j.** Quantification of the immunoblot intensities of PTPRT and its fragments, pSTAT3^Y705^ shown in panel i). Error bars denote S.E.M. (n= 3 independent experiments; *, *P* < 0.001). **k.** Protein extracts from primary cortical neurons of wild type with infection of lentiviral shRNAs at indicated DIV were assayed by Western blot for the presence of PTPRT and its fragments, and pSTAT3^Y705^. **l-m.** Quantification of the immunoblot intensities of PTPRT and its fragments, pSTAT3^Y705^ shown in panel k). Error bars denote S.E.M. (n = 3 independent experiments; *, *P* < 0.001).

pSTAT3^Y705^ is a well characterized substrate of PTPRT in cancer cells. While *Ptprt* is preferentially expressed in the central nervous system, thus it is worth investigating whether PTPRT dephosphorylates pSTAT3^Y705^ in the brain, especially in neurons. First, we examined the dynamics of PTPRT in the developmental mouse brain. Indeed, the full length of PTPRT showed an increasing trend with the postnatal age (Fig. 1i-j). On the contrary, the level of pSTAT3^Y705^ presented a decreasing tendency. Unlike full length, PTPRT-CTF2, the product of ADAM10 cleavage, also decreases gradually during the postnatal stage, showing efficient cleavage of PTPRT-CTF2 by γ-secretase with the maturing of the brain. It seems that a low level of PTPRT may be necessary for maintaining a relatively high level of pSTAT3^Y705^, and γ-secretase mediated processing may be involved in it. A high level of pSTAT3^Y705^ is essential for neurogenesis and development in the early stage, while minimum pSTAT3^Y705^ is needed for adults. Next, to examine whether PTPRT directly dephosphorylates pSTAT3^Y705^, using shRNAs against *Ptprt* mRNA we knocked down its expression in wild-type primary cortical neurons. As expected, knockdown of *Ptprt* significantly led to an increase in the level of pSTAT3^Y705^ (Fig. 1k-m), suggesting a crucial role of PTPRT in the dephosphorylation of pSTAT3^Y705^ in neurons.

### Membrane to nuclear translocation of PTPRT intracellular domain dephosphorylates STAT3

Nuclear translocation of the released intracellular domain by γ-secretase plays an important role in the regulation of intramembrane proteolysis-mediated signaling. To examine whether a nuclear localization signal (NLS) in PTPRT is responsible for the nuclear translocation of its intracellular domain, using several different types of online software we analyzed and identified that there was putative NLS (KRRKLAKKQK) at the N terminal of PICD with a high score (http://mleg.cse.sc.edu/seqNLS)[35]. To test whether the released PICD could translocate to the nucleus via the leading of NLS or not, we cloned PICD^wt^ and PICD^deltaNLS^ (without NLS) sequence to a vector fused with mCherry, and then transfected them to HEK293 cells respectively. Unlike mCherry with whole-cell dispersion, PICD^wt^-mCherry is specifically distributed in the nucleus (Fig. 2a). The PICD^wt^ also completely localized in the nucleus of live N2a cells (Fig.2b). However, the deletion of the putative NLS led to little accumulation of PICD in the nucleus. Due to rapid degradation by the proteasome, it’s hard to demonstrate the presence of PICD in the nucleus directly. As a result, we employed a Gal4-UAS based transcription activation system to confirm the nuclear localization of PICD (Figure S2c-d). The system showed that activation of α-secretase shedding by treatment of PMA increased the transcriptional signaling by more PICD nuclear translocation, while DAPT treatment decreased the PICD mediated nuclear transactivation. The evidence showed that the endogenous PICD could also translocate to the nucleus.

**Fig. 2.**
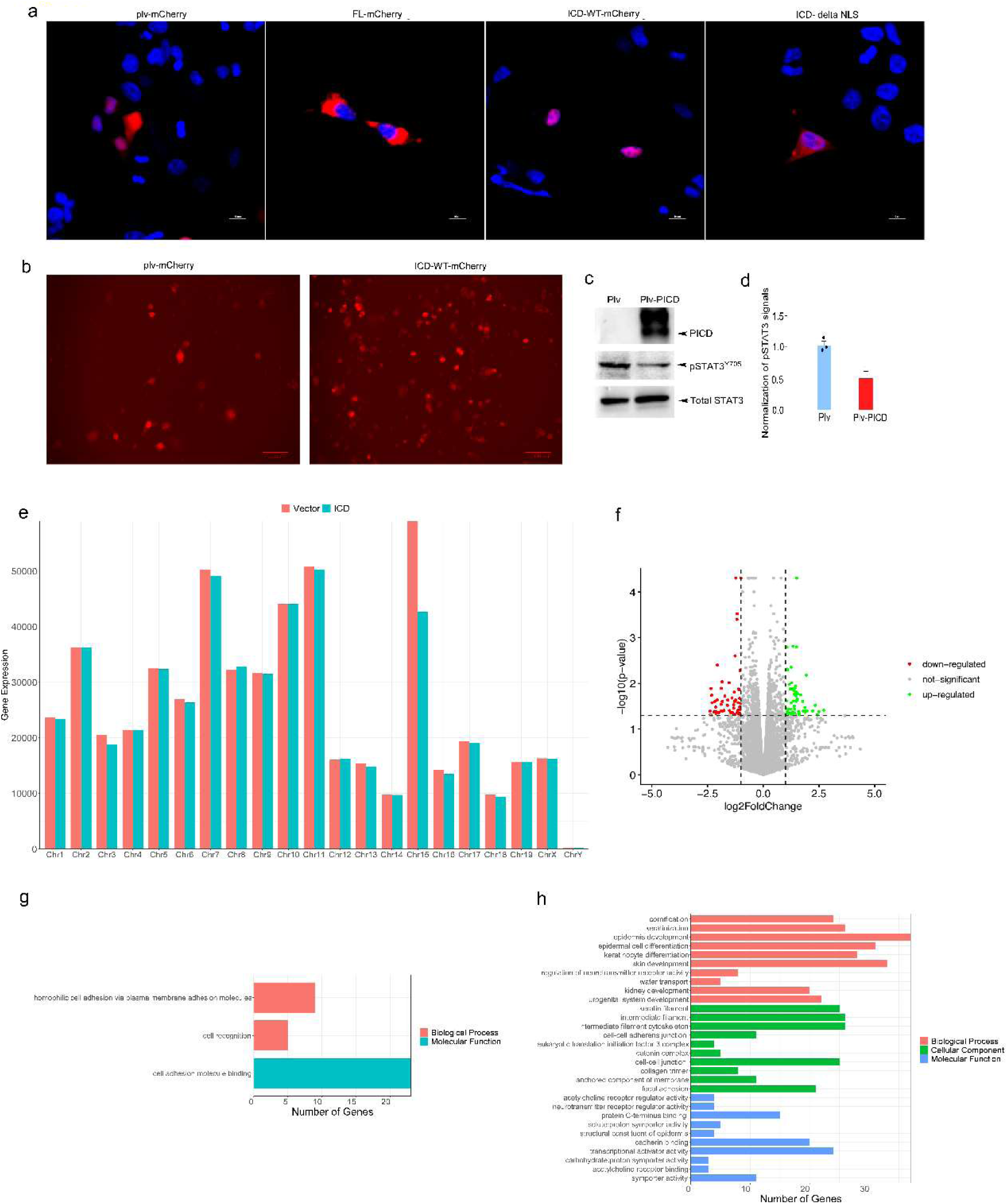
Nuclear localization of PICD decreases pSTAT3^Y705^ and regulates gene expression in neural cells. **a.** Representative figures show the specific nuclear localization of PICD led by its NLS in Plv, PTPRT-, PICD- and PICDdetaNLS-mcherry expressed HEK293 cells. **b.** Representative figures show the nuclear localization of PICD in live N2a cells. **c.** Protein extracts from Plv- and PICD-mCherry N2a cells were assayed by Western blot for the presence of PICD and pSTAT3Y705. **d.** Quantification of the immunoblot intensities of pSTAT3Y705 shown in panel c). Error bars denote S.E.M. (n= 3 independent experiments; *, *P* < 0.005). **e.** The sum of expression of all genes located in each chromosome. **f.** The genes with significantly different expression between the control vector and PICD groups, red points represent down-regulated genes in the PICD group, green points represent up-regulated genes in the PICD group, grey points represent little significant difference. **g.** The GO annotation contains the *Smagp* gene on chromosome 15. **h.** The number genes annotated to top 10 biological processes, cellular component, and molecular functions, respectively, on chromosome 15.

The PTPRT intracellular fragment contains two phosphatase domains, so it was very interesting to investigate whether the released ICD after γ-secretase cleavage still takes the dephosphorylating activity. The phosphorylated STAT3 at Y705 would form a dimer and translocate to the nucleus and then acts as a transcriptional factor. We then transfected N2a cells with PICD, and our data demonstrated that PICD significantly decreased the level of pSTAT3^Y705^ in N2a cells (Fig. 2c-d).

### PICD regulates gene expression

Sequential processing and then releasing the signaling intracellular domain has been described for several RIP participating type 1 transmembrane proteins, however the mechanism of PTPRT signaling has not been delineated yet. We then hypothesized that nuclear localized PICD could take some transcription factor like function, such as NICD. At least, PICD could regulate gene transcription indirectly by dephosphorylating pSTAT3^Y705^. To test this hypothesis, we generated inducible isogenic N2a cell lines stably expressing the mCherry-tagged PICD as well as mCherry control and subsequently performed RNA deep sequencing (RNA-seq). We then determined the global transcriptome changes of cells expressing the PICD versus cells expressing mCherry as control. We compute the sum of expression of all genes located in each chromosome. And the differences in gene expression between the control vector and the ICD group were presented (Fig. 2e). If Delta > 0, the sum of gene expression is up-regulated, otherwise, the sum of gene expression is down-regulated. The expression of chromosome 15 has the largest down-regulated expression between the control vector and the ICD group (-27.7%). The largest differential expression location of chromosome 15 is NC_000081.6:47539988-47540137, which can not be annotated to a defined gene. This location contributes to the main difference in chromosome 15 (the delta expression of chromosome 15 is 42659.0-58997.5=-16338.5, the delta expression of NC_000081.6:47539988-47540137 is 16273-32655=-16382). The significant difference between genes was defined with *p*-value <0.05 and the absolute value of log2Foldchange more than 1 (Fig. 2f). Red points represent significantly down-regulated genes with log2Foldchange < -1, green points represent significantly up-regulated genes with log2Foldchange >1. The grey points are genes without significant difference between the control vector and the ICD group. Down-regulated and up-regulated genes by PICD expression were presented in the heatmap (Figure S2c-d). The *Ptprt* gene is significantly up-regulated in PICD group. There are 128 genes with a significant difference between the control vector and PICD, which contains 62 down-regulated and 66 up-regulated genes.

Since the *Ptprt* gene is located on chromosome 2. We selected the biological process and molecular functions that contain the *Ptprt* gene (Figure S2a, Figure S2e). All genes on chromosome 2 to the GO database and KEGG database were then annotated (Fig. 1b, Figure S3f). Indeed, most of those biological processes and molecular functions are related to dephosphorylation, phosphatase, or cell adhesion. The annotation of the *Ptptr* gene in the KEGG database is “receptor-type tyrosine-protein phosphatase T”. There are no genes that have the same KEGG annotation as the *Ptprt* gene. We then annotated all genes on 21 chromosomes (19+XY) to GO and KEGG databases. There are no genes that have the same KEGG annotation as the *Ptprt* gene too. But 978 genes have the same GO annotations with the *Ptprt* gene. Amongst, we identified that *Smagp* (small cell adhesion glycoprotein)[36, 37] located on chromosome 15 showedhas the largest differential expression between the control vector and the ICD group. We then selected the biological process and molecular functions that contain *Smagp* gene (Fig. 2g, Figure S3e). All genes on chromosome 15 to the GO database and KEGG database were annotated (Fig. 2h, Figure S3h). Next, to validate the effect of PICD on targeted gene mRNA expression, levels of downregulated and upregulated candidates *Smagp*, *Shisal2a*, *Pax4*, *Polr3gl*, *Ptprtp*, *Mir6379*, and*cye* mRNA expression were examined (Figure S3a-b). Indeed, qPCR data is consistent with RNAseq analysis.

### The level of PTPRT expression decreased in the brains of human AD and model mice

PTPRT is highly expressed in the mammalian brain and regulates STAT3 phosphorylation. Its function in neurodevelopment has been reported. To investigate whether and how alteration in PTPRT signaling occurs in age-related neurodegenerative diseases such as AD remains unclear, first, the RNA-seq data set of human AD brain samples were downloaded from NCBI and analyzed. Surprisingly, *Ptprt* mRNA expression was significantly downregulated in several brain regions of human AD including PFC, HP, EC, and TC compared to that of controls (Fig. 3a). Next, using immunohistochemistry (IHC) the levels of PTPRT in age- and sex-matched postmortem human control and AD brain samples were observed. Indeed, a significant decrease in the intensities of PTPRT signals was found in the PFC and hippocampus of AD compared to age-matched control samples (Fig. 3b-c). Western blot data from human AD brain tissues further confirmed a decrease in the protein level of PTPRT full length and two CTF cleavages (Fig. 3d-e).

**Fig 3.**
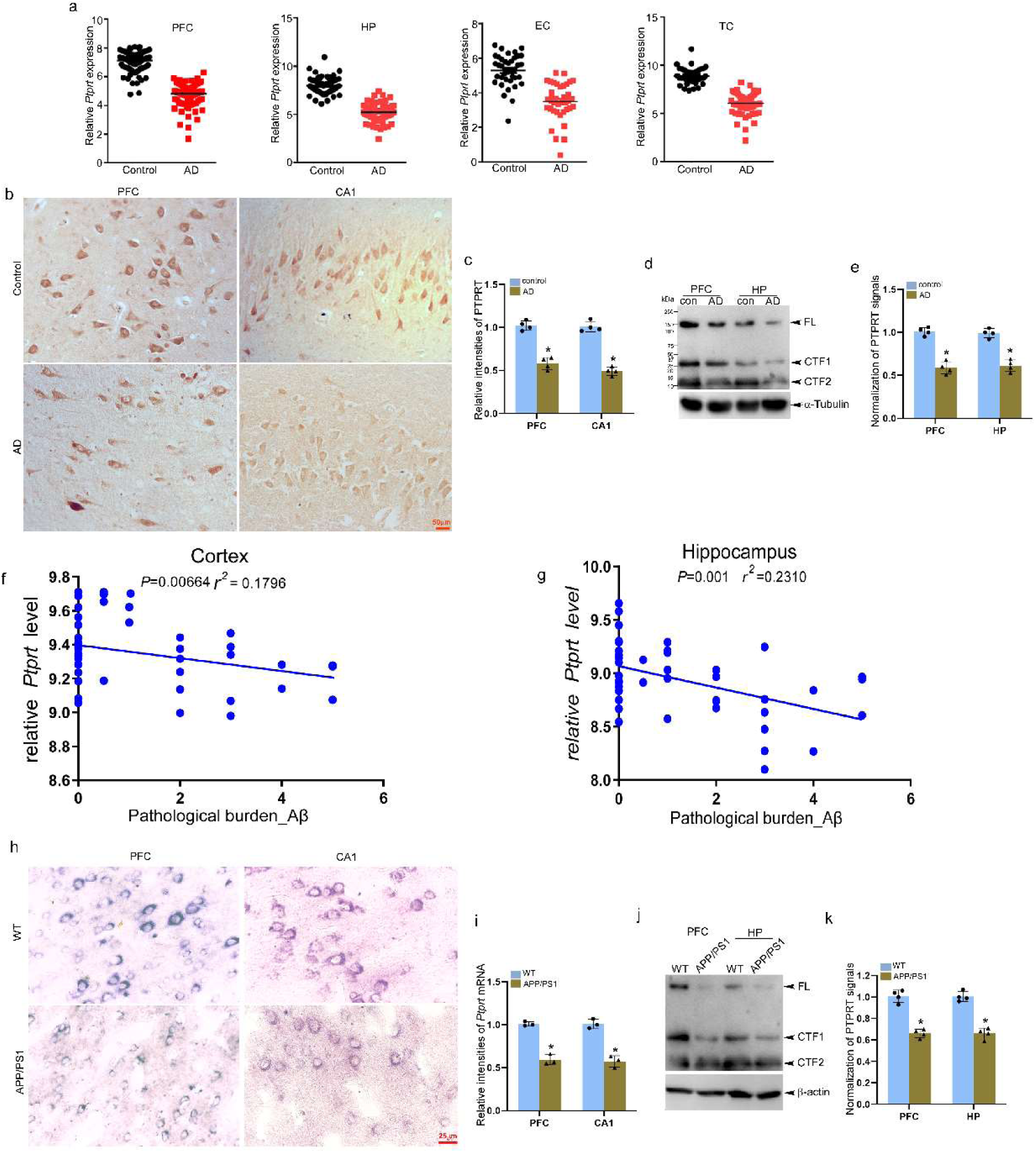
mRNA and protein levels of PTPRT decreased in the brains of human AD and model mice. **a.** The bioinformatic analysis shows the downregulation of *Ptprt* expression in the brains of AD patients. **b.** 20 μm paraffin sections of PFC and hippocampus of control and AD patients were immunostained with PTPRT (Brown) antibody. Scale bar, 50μm. **c.** Quantification of normalized intensities of PTPRT illustrated in b). Error bars denote S.E.M. (n= 4 patients per group; *, *P* < 0.001). **d.** Protein extracts from frozen postmortem PFC and hippocampal tissues of control and AD patients were assayed by Western blot for the presence of PTPRT and its fragments. **e.** Quantification of the immunoblot intensities of PTPRT shown in panel d). Error bars denote S.E.M. (n= 4 patients per group; *, *P* < 0.001). **f-g.** The bioinformatic analysis shows the significantly negative correlation between *Ptprt* expression and pathological amyloid β burden in the brains of APP/PS mice. **h.** 10 μm cryostat brain sections from 12-month-old wild type and APP/PS mice were stained by ISH. Scale bar, 50μm. **i.** Quantification of normalized intensities of PTPRT illustrated in h). Error bars denote S.E.M. (n= 3mice per genotype; *, *P* < 0.001). **j.** Protein extracts from fresh PFC and hippocampal tissues of 12-month-old wild type and APP/PS mice were assayed by Western blot for the presence of PTPRT and its fragments. **k.** Quantification of the immunoblot intensities of PTPRT shown in panel j). Error bars denote S.E.M. (n= 3mice per genotype; *, *P* < 0.001).

Next, to assess whether AD mouse models can recapitulate decreased mRNA expression and protein level of PTPRT found in human AD brain, first, RNA-seq data from APP/PS1 mouse brain samples were downloaded from NCBI, and the expression level of *Ptprt* and the Aβ pathological burden in the brain of APP/PS1 mice were quantified. We found that the level of *Ptprt* expression showed a significant negative correlation with β-amyloid pathological burden in both cortex (r^2^ = 0.1796, *P* = 0.0064) and) and hippocampus (r^2^ = 0.2310, *P* = 0.001) of APP/PS1 mice (Fig. 3f-g). Next, using in situ hybridization and Western blot we further verified the decrease in the levels of PTPRT in PFC and CA1 of 12-month-old APP/PS1 mice (Fig. 3h-k).

Lastly, we detected the level of by pSTAT3^Y705^ immunostaining the brain of AD patients and mouse models. As was expected, the intensities of pSTAT3^Y705^ were significantly upregulated in PFC and hippocampal CA1 of human AD and APP/PS mouse brains compared to control groups (Fig. 4a-d). In addition to decreased *Ptprt* expression, there may be some other factors that contribute to the nuclear accumulation of pSTAT3^Y705^.

**Fig 4.**
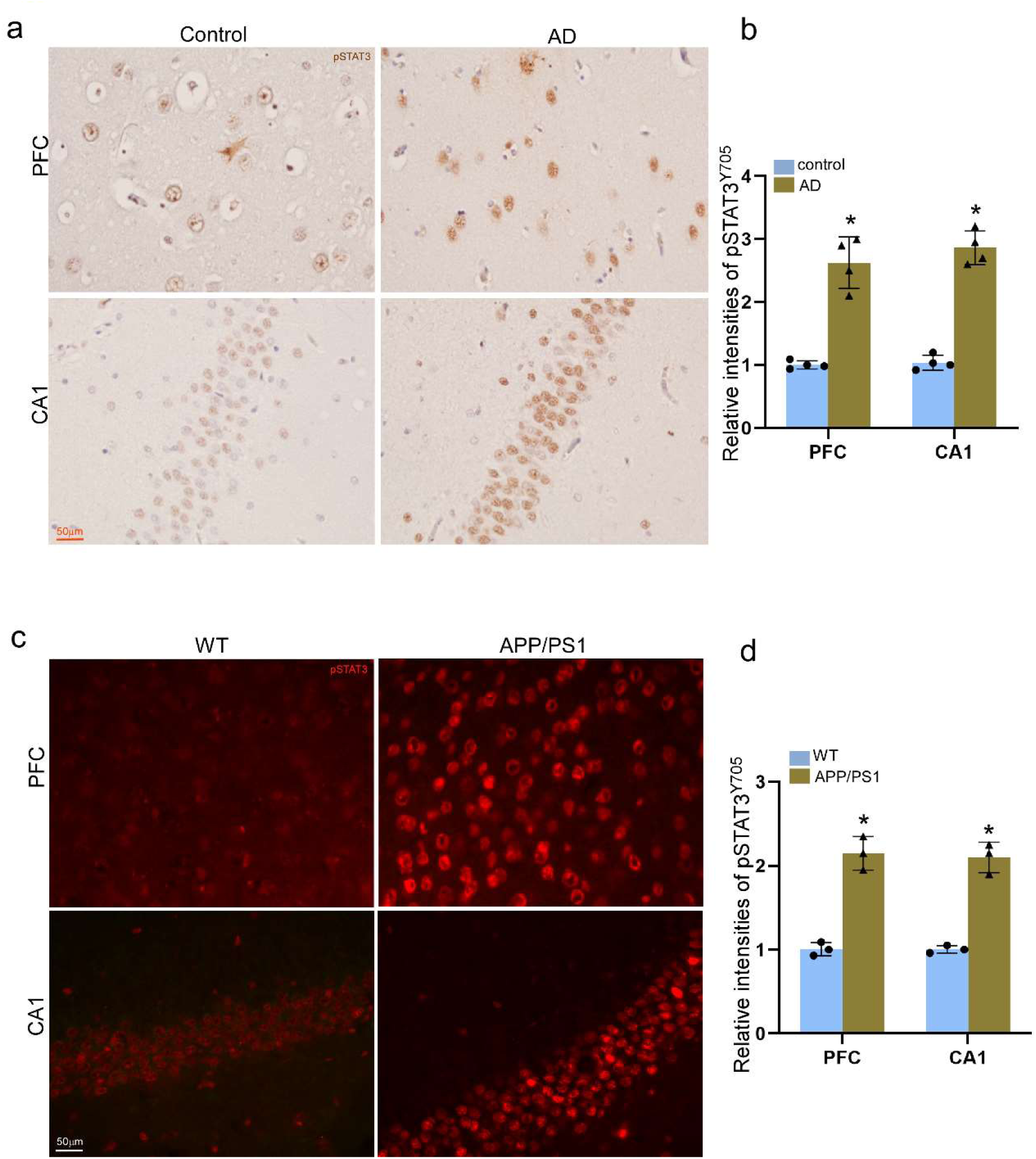
Increased pSTAT3^Y705^ found in both human AD and APP/PS mouse brains. **a.** 20 μm paraffin sections of PFC and hippocampus of control and AD patients were immunostained with pSTAT3^Y705^ (Brown) antibody. Scale bar, 100μm. **b.** Quantification of normalized intensities of pSTAT3^Y705^ illustrated in a). Error bars denote S.E.M. (n= 4 patients per group; *, *P* < 0.001). **c.** 10 μm cryostat brain sections from 12-month-old wild type and APP/PS mice were immunostained with pSTAT3^Y705^ (Red) antibody. Scale bar, 50μm. **d.** Quantification of normalized intensities of pSTAT3^Y705^ illustrated in c). Error bars denote S.E.M. (n= 3mice per genotype; *, *P* < 0.001).

### PICD prevents Aβ deposition and pathology both *in vitro* and *in vivo*

AAV delivery of the *Ptprt* gene to the brain could be an ideal way for *in vivo* gene therapeutic intentions. However, the full length of *Ptprt-*coding sequence is about 4.4 kb, together with the promoter, ITR, and other regulatory sequences the whole size will exceed the maximum accommodation. PICD with half of the full size makes it a good fit for AAV-packaging. Using microinjection of viral particles containing PICD into the hippocampus of 9-month-old mice we observed its effect on AD pathologies *in vivo*. The efficiency of PICD viral infection was firstly verified by immunohistochemistry and Western blot analysis. Notably, PICD significantly decreased pSTAT3^Y705^ levels in the hippocampus of APP/PS1 mice (Fig.S5).

Next, to assess whether increased pSTAT3^Y705^ induced by loss of the intracellular domain of PTPRT contributes to Alzheimer’s pathogenesis as well as neurodegeneration of AD, we decided to examine the effect of the intracellular domain of PTPRT on Aβ deposition *in vitro* and *in vivo*. First, primary cortical neurons from wild type and APP/PS1 mouse embryos (E16.5) were infected with lentiviral PICD at DIV 5. Neurons were collected at DIV 14, and immunostained with pSTAT3^Y705^ and 6E10 and MOAB2 antibodies. Indeed, PICD-expressing neurons showed a significant decrease in pSTAT3^Y705^ and 6E10 signals (Fig. 5a-d).

**Fig. 5.**
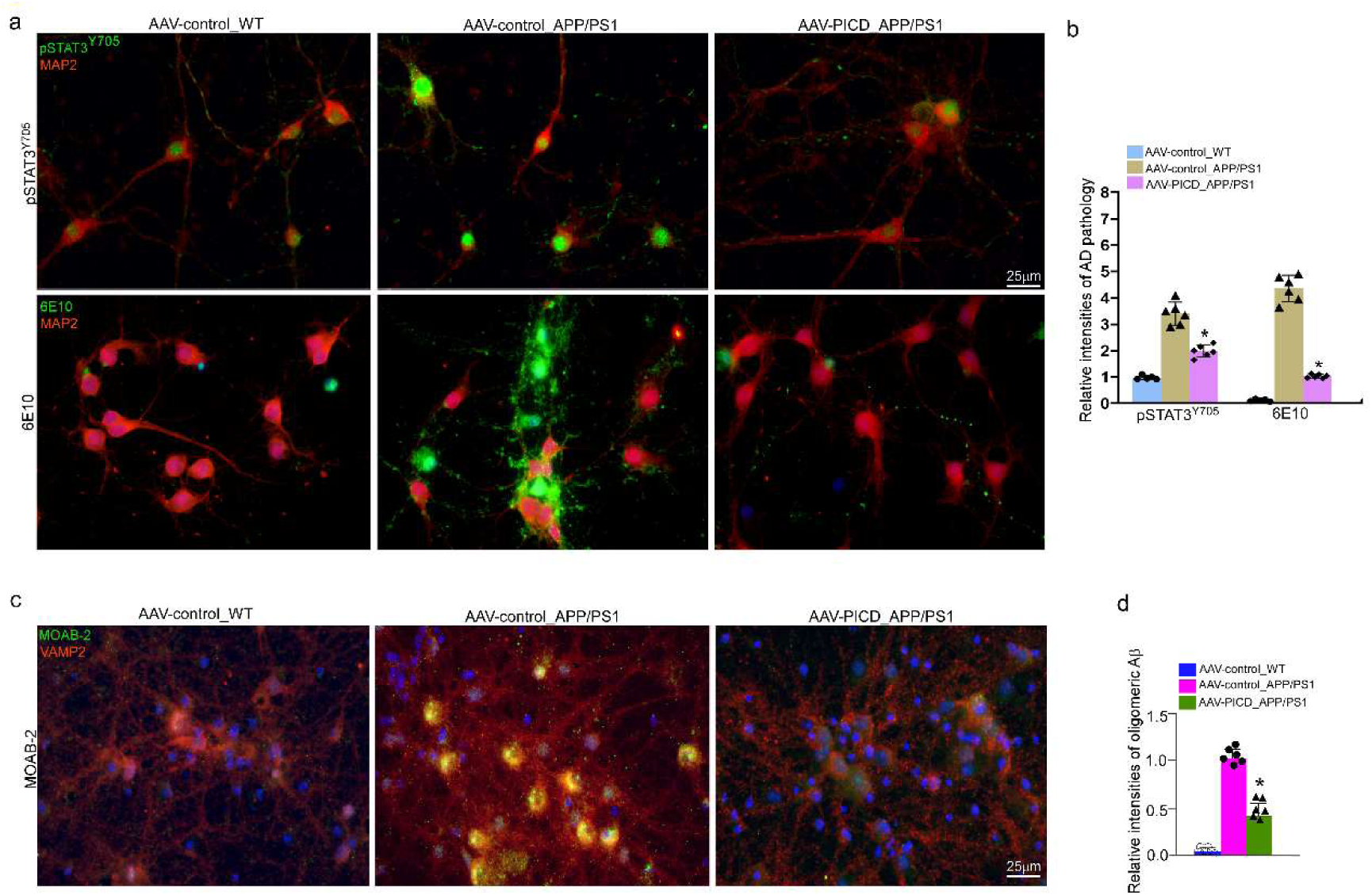
Overexpression of PICD prevents Aβ production in APP/PS1 neurons. **a.** Representative IF images showing the levels of pSTAT3Y705 and 6E10 in cortical neurons of wild type and APP/PS1 mice. **b.** Quantification of intensities of pSTAT3 and 6E10 immunoreactivities in the hippocampus normalized to an area in square millimeters. Data are presented as mean ± S.E.M. (*, P < 0.005) compared with viral vector-injected wild type mice as measured by Student’s t-test (n= 6 independent E16.5 primary cultures per genotype). **c.** Representative IF images showing the levels of oligomeric Aβ in cortical neurons of wild type and APP/PS1 mice. **d.** Quantification of intensities of MOAB-2 immunoreactivities in the hippocampus normalized to an area in square millimeters. Data are presented as mean ± S.E.M. (*, P < 0.005) compared with viral vector-injected wild type mice as measured by Student’s t-test (n= 6 independent E16.5 primary cultures per genotype).

Next, using the histological and Western blot we examined the effect of overexpression of PICD on Aβ deposition and pathology. As expect, the induction of PICD led to a significantly reduced pathological amyloid β burden (Fig. 6a-d). Western blot results showed decreased cleaved caspase 3 in the hippocampus of APP/PS1 mice compared to controls of the viral vector (Fig.6e-f).

**Fig. 6.**
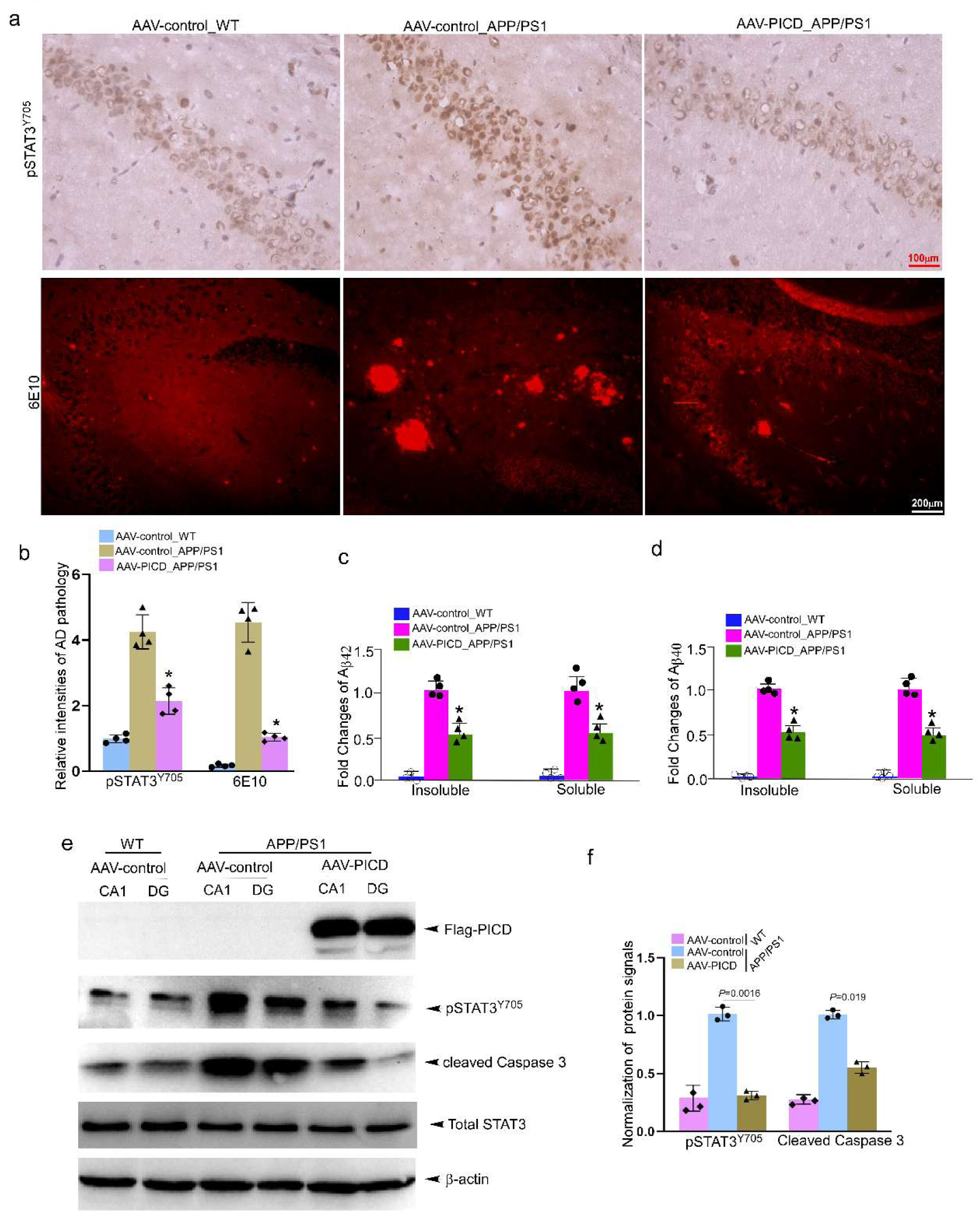
PICD leads to decreases in the levels of pSTAT3 and Aβ deposition. **a.** Representative IHC images showing the levels of pSTAT3^Y705^ and 6E10 in the hippocampus of wild type and APP/PS1 mice. The viral particles of AAV-control or AAV-PICD was injected into the hippocampal regions of 9-month-old mice, and cryostat slices were either immunostained with pSTAT3^Y705^ and 6E10 antibodies at 12 months. Bars were labeled as indicated. **b.** Quantification of intensities of pSTAT3^Y705^ and 6E10 immunoreactivities in the hippocampus normalized to an area in square millimeters. *, *P* < 0.001 compared with viral vector-injected wild type mice as measured by Student’s *t-*test. (WT, AAV-control: n=4; APP/PS1, AAV-control, n=4; APP/PS1, AAV-PTPRT-ICD, n=4). **c.** ELISA analysis shows a significant reduction in soluble Aβ_42_, and a trend toward a reduction in soluble oligomeric Aβ in 12-month-old APP/PS1 mice with overexpression of PICD. **d.** ELISAs on brain lysates were performed using MOAB-2 as the capture antibody and MOAB-2- and Aβ_42-_biotin as detection antibodies. *, *P* < 0.005 (n = 4 mice per genotype). **e.** Protein extracts from fresh hippocampal CA1 and DG tissues of 12-month-old wild type and microinjection of AAV-control or AAV-PICD were assayed by Western blot for the presence of PICD, pSTAT3^Y705^, and cleaved Caspase-3. β-actin was loading control. **f.** Quantification of the immunobloted intensities of pSTAT3^Y705^ and cleaved Caspase-3 shown in panel e). Error bars denote S.E.M. (n = 3 mice per genotypes; *, *P* < 0.001).

### PICD prevents synaptic dysfunction and behavioral deficits in APP/PS1 mice

To investigate whether the effect of PICD on the basal synaptic transmission, we generated input/output (I/O) curves by measuring field-excitatory by stimulation of the Schaffer collaterals at increasing stimulus intensities at 12-month-old mice. APP/PS1 mice with microinjection of PICD viral-like particles into the hippocampus exhibited bigger fEPSP slopes and amplitudes at all stimulus intensities tested and had significantly increased maximum fEPSPs relative to wild type mice (Fig. 7a). The deficits of long-term potentiation (LTP) in AD mice have been well documented before[38, 39]. We, therefore, investigated LTP in12-month-old wild type and APP/PS1 mice. Due to impaired LTP in the APP/PS1 mice, and the smaller absolute fEPSP may account for the LTP deficits. Next, we increased the stimulus intensity of the APP/PS1 mice to match baseline fEPSP magnitudes to those of wild type mice. The stimulus intensity used to elicit LTP in the wild type sections was approximately 30% of the maximum fEPSP slopes, equating to a value of ~0.45mV/ms, which is below the max value of the wild type mice. LTP magnitudes in these experiments did not significantly differ in the percentage of potentiation when these stronger stimulus intensities were used, indicating that reduced basal transmission does not likely account for the deficits in LTP in the wild type mice. As with the APP/PS1 mice, raising baseline fEPSPs to wild type levels did not result in significantly different LTP, and thus, the data were pooled. Meanwhile, we also lowered the baseline fEPSP of the wild type mice to match APP/PS1 mice and found no difference in potentiation as compared with the normal LTP protocol. All the data were pooled from the APP/PS1 experiments, where baseline fEPSPs were matched to wild type levels, with the experiments where the normal LTP protocol was used. We also pooled the data from the wild type experiments, where baseline fEPSPs were lowered to APP/PS1 levels, with data from the normal LTP protocol. In consistent with reports from other studies, our pool data showed that the reduced LTP in the APP/PS1 as compared to wild-type mice. Indeed, LTP in the APP/PS1 mice with injected PICD viral-like particles was significantly improved despite also having weaker fEPSPs relative to wild type controls (Fig. 7b).

**Fig 7.**
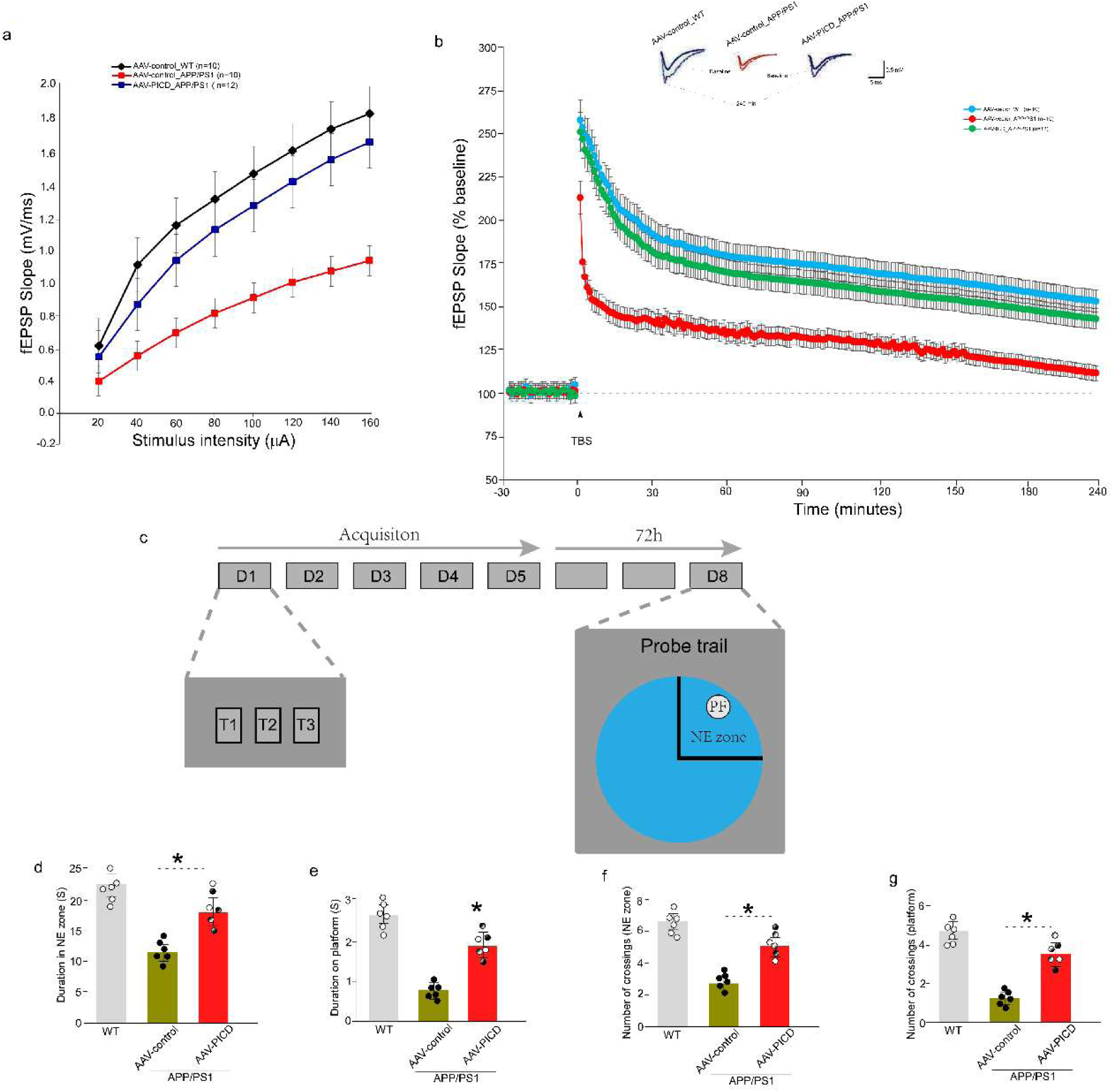
PICD improves synaptic function and working memory in APP/PS1 mice. **a.** Input/output (I/O) curves and representative fEPSPs at increasing stimulus strengths are shown for WT, APP/PS1 mice with injected viral vector, PICD, respectively. Bigger fEPSPs are evoked in 12-month-old APP/PS1 mice with microinjection of the viral PICD particles compared to the control viral particles, indicating improvement of impaired synaptic transmission. WT, viral control, 1.43mV/ms ± 0.16 mV/ms, n = 8 slices from 4 mice (10/5); APP/PS1, viral control 0.51mV/ms ± 0.08mV/ms, n = 10/5, *P* <0.001; APP/PS1, PICD, 1.26mV/ms ± 0.13mV/ms, n = 12/6, *P* < 0.001. **b.** fEPSP slopes were recorded and were expressed as the percentage of the pretetanus baseline. Representative fEPSPs before (solid line) and 60 min after the induction of LTP (dotted line) are shown. Impaired LTP in APP/PS1 mice was markedly improved in the overexpression of human PICD catalytic domains. The amount of potentiation of fEPSPs between 0 and 10 min after HFS was 143% ±15% in AAV control-expressing APP/PS1 mice (n = 10/5), and was significantly improved in PICD-expressing APP/PS1 mice (235% ± 16%, n = 12/6, *P* < 0.001), compared to WT mice (256% ±19%, n = 10/5, *P* =0.0058). The amount of potentiation between 50 to 60 min after HFS was 229% ± 16% in PICD-expressing APP/PS1 mice and was not significantly different from WT mice (235% ± 18%, *P* = 0.167), compared to AAV control expressing APP/PS1 mice (129% ± 10%). Significance was determined using a two-way ANOVA. **c.** Pipeline of the Morris water maze experiment. Training consisted of daily sessions (three trials per session) during 5 consecutive days. Mean inter-trial interval was 15 min. 72 h after the last training trial (day 8), retention was assessed during probe trial in which the platform was no longer present. **d-g.** PICD alone leads to improvement of spatial learning and memory in APP/PS1 mice. (c-d) Duration in NE zone and the submerged platform area in the probe test. Data were expressed as mean ± **SEM** (**n** = 6/group). \**p* < 0.05, compared with control. (e-f) Number of crossing the NE zone and submerged platform area times in the probe test. Data were expressed as mean ± **SEM** (**n** = 6/group). \**p* <0 .01, compared with control.

Next, the learning and spatial memory was further tested using the Morris water maze (Fig.7c). The test was conducted in wild type and APP/PS1 mice after one month of microinjection of AAV-PICD into the hippocampus at the age of 9 months. In contrast to wild type mice, the total time of exploration in the target quadrant shows that the swimming time of APP/PS1 mice in the target quadrant is significantly reduced (*P* = 0.0366, n = 6 mice). Nevertheless, PICD is overexpressed group can increase the swimming time of APP/PS1 mice in the target quadrant (*P* = 0.0069, n = 6 mice) (Fig. 7d-e). Meanwhile, while the latency of APP/PS1 mice crossing the platform in AAV-control group was significantly increased (*P* = 0.0367, n = 6 mice), the latency of mice crossing the platform in the overexpression PICD group was significantly reduced (*P* = 0.0236, n = 6 mice) (Fig. 7f-g). This date suggests that the induction of PICD in the hippocampus of APP/PS1 mice can significantly restore the learning ability and spatial learning memory compared to the control groups.

## Discussion

γ-secretase mediated RIP plays an essential role in several biological processes[2]. In this study, the type 1 transmembrane protein PTPRT was identified as a novel substrate of ADAM10 and presenilin 1/γ-secretase, and preferentially restricted to the brain and lessened neurodegeneration of AD. PTPRT was cleaved by presenilin 1/γ-secretase after the ADAM10 mediated shedding, released the functional domain PICD. The nuclear translocation of PICD efficiently dephosphorylates pSTAT3 in the nucleus. Moreover, the decreased PTPRT found in the brains of both human AD and APP/PS1 mouse brains led to an excessive nuclear accumulation of pSTAT3. PICD alone not only decreased the pSTAT3 level and Aβ deposition, but also could improve behavioral deficits in APP/PS1 mice.

STAT3 has been identified as a substrate of PTPRT. By dephosphorylating STAT3 at Y705, PTPRT blocks dimerization and nuclear translocation of STAT3 and subsequently inhibits tumorigenesis[22]. AS a result, PTPRT has become a potential therapeutic target for certain cancers. As a central nervous system-specific protein, PTPRT may have a similar function in the adult brain. Our findings show that PTPRT could wipe phosphate at STAT3 in neurons and it may be a key factor in regulating phosphorylation of STAT3 during brain development. A high level of pSTAT3 is necessary for motor neuron differentiation during the embryonic stage when the *Ptprt* shows a lower expression level in our result. In the adult mouse brain, high expression of *Ptprt* matches lower pSTAT3, which indicates the pivotal role of PTPRT in downregulating STAT3 activation.

It has been hypothesized that rapid activation of STAT3 following acute injuries is neuroprotective due to the subsequent transcriptional upregulation of neurotrophic genes[40–42]. However, chronic activation of STAT3 is detrimental, especially in neurodegenerative diseases. Microglial JAK/STAT3 pathway activation was found in the brain of Parkinson’s disease, which would initiate the inflammatory responses in the brain and lead to neurodegeneration and neural death ultimately[43, 44]. STAT3 activation triggers astrogliosis in Alzheimer’s disease[45], and the deletion of STAT3 in astrocyte would reduce Aβ plaque burden and improve the cognitive performance in APP/PS1 mouse model[46]. Aβ induced accumulation of pSTAT3 in neurons directly activated the apoptotic pathway. The Aβ-pSTAT3-Aβ cycle may play a pivotal role in the progressive process. Ser727 at STAT3 is another important phosphorylation site which also involved in the STAT3 activation. Previous work showed that the phosphorylation of STAT3 at Ser727 would induce the dimer formation, which subsequently led to the phosphorylation of STAT3 (Y705). The nuclear translocation of dimerized pSTAT3^Y705^ upregulated the expression of BACE1 and then promoted the generation of amyloid beta[47]. The inhibition of STAT3 activity may be helpful to reduce the production of b-amyloid and slow the progression. Therefore, inhibitions of the JAK/STAT3 pathway become potential therapeutics for neurodegenerative diseases. However, the risk of adverse effects by nonselective inhibition of STAT3 activity in the whole body still deserved serious consideration. Failure to cross the blood-brain barrier (BBB) is also an important concern for many molecules[48].

Pathological alterations in postsynaptic density composition, glutamatergic synaptic transmission, and dendritic spine are complicated in the process of Alzheimer’s disease[49]. In the late stage, the significant loss of neurons exacerbates the situation. Caspase 3 is not only a final executor of apoptotic cell death but also a participator in synaptic dysfunction via a non-apoptotic dependent manner[50–52]. Increased caspase 3 activity in spines mediates GluR1 removal from postsynaptic sites in the hippocampal synapses of Tg2576 mice[53]. In this work, microinjection of AVV-PICD into the hippocampus of APP/PS1 mice significantly reduced the levels of cleaved caspase 3, though the number of hippocampus neurons was not significantly changed. Overexpression of PICD not only restores the LTP impairment but also improves working memory in APP/PS1 mice. In addition, the changed expression of neuronal circuit associated genes by PICD may also contribute to the rescue of LTP and improvement of working memory. Though lots of evidence shows that the blocking of STAT3 signal could stimulate synaptogenesis[54, 55], it’s still worth investigating whether the nuclear PICD can directly bind to genomic DNA and sever as a transcriptional regulator directly in the future.

The pathogenesis of Alzheimer’s disease is complex, and our understanding of the mechanism still needs to be improved. In the past two decades, therapeutic effects against AD mainly focused on β-amyloid-producing γ-secretase, and some of the potent γ-secretase inhibitors (GSIs) were found. However, up to now none of these potential drugs were successful in clinical trials. One of the most important reasons is that those GSIs would non-selectively block the cleavage of many other substrates[56], some of which may have essential physiological roles[57]. Once blocked, serious adverse effects would happen, like squamous cell carcinoma[58]. In this study, we found that abnormal ADAM10- and presenilin1/γ-secretase-mediated proteolysis of PTPRT is involved in the pathological accumulation of pSTAT3 in the nucleus of neurons in AD brain, and demonstrated that PICD significantly improves neuropathological changes and behavioral deficits in APP/PS1 mice, indicating a potentially novel therapeutic strategy for AD.

## Acknowledgments

We are grateful to Bart De Strooper from the KU Leuven in Belgium for granting Ps1/Ps2 double knockout MEF cells, and Yufang Zheng from Fudan University for granting ADAM10^-/-^ and ADAM17^-/-^ HEK293 cells. We also thank Prof. Juming Zhou (Kunming Institute of Zoology) for providing us the constructs for GAL4/UAS System. The authors also thank Dr. Karl Herrup for critical reading the manuscript and constructive suggestion. This work was financially supported by the National Science Foundation of China (91649119 and 92049105) to J.L.; and the Peking University (BMU2019YJ001) to J.L.; and the key basic project of Yunnan Province (E039030401) to J.L.

## Contributions

S.L. and J.L. conceived and designed all experiments. S.L., Z.Z., and L.Y. conducted the majority of experiments including Immunohistochemical analysis, western blot, confocal microscopy, fluorescence assay, electrophysiological and animal assays. Z.Z. contributed in the brain slices recordings. L.L., Z.M., and J.L. performed RNA-seq and data analysis; S.L., Z.Z., and L.Y. performed primary cultures, viral particle preparation and infection, and viral microinjection; S.L., and J. L. drafted the manuscript with input from all other authors. All authors read and approved the final manuscript.

