## Supplementary Information for "ADAM10- and presenilin 1/γ-secretase-dependent cleavage of PTPRT mitigates neurodegeneration of Alzheimer’s disease"

#### Table S1. Sequence of shRNAs against *Ptprt* mRNA

| *gene* | *Target sequence (Blue)* |
|---|---|
| scramble | ccgggcgcgatagcgctaataatttctcgagaaattattagctatcgcgctttttg |
| mouse *ptprt* | ccggcctcatttctatcaggtgatactcgagtatcacctgatagaaatgaggtttttg |

### Supplementary materials

**Figure S1**

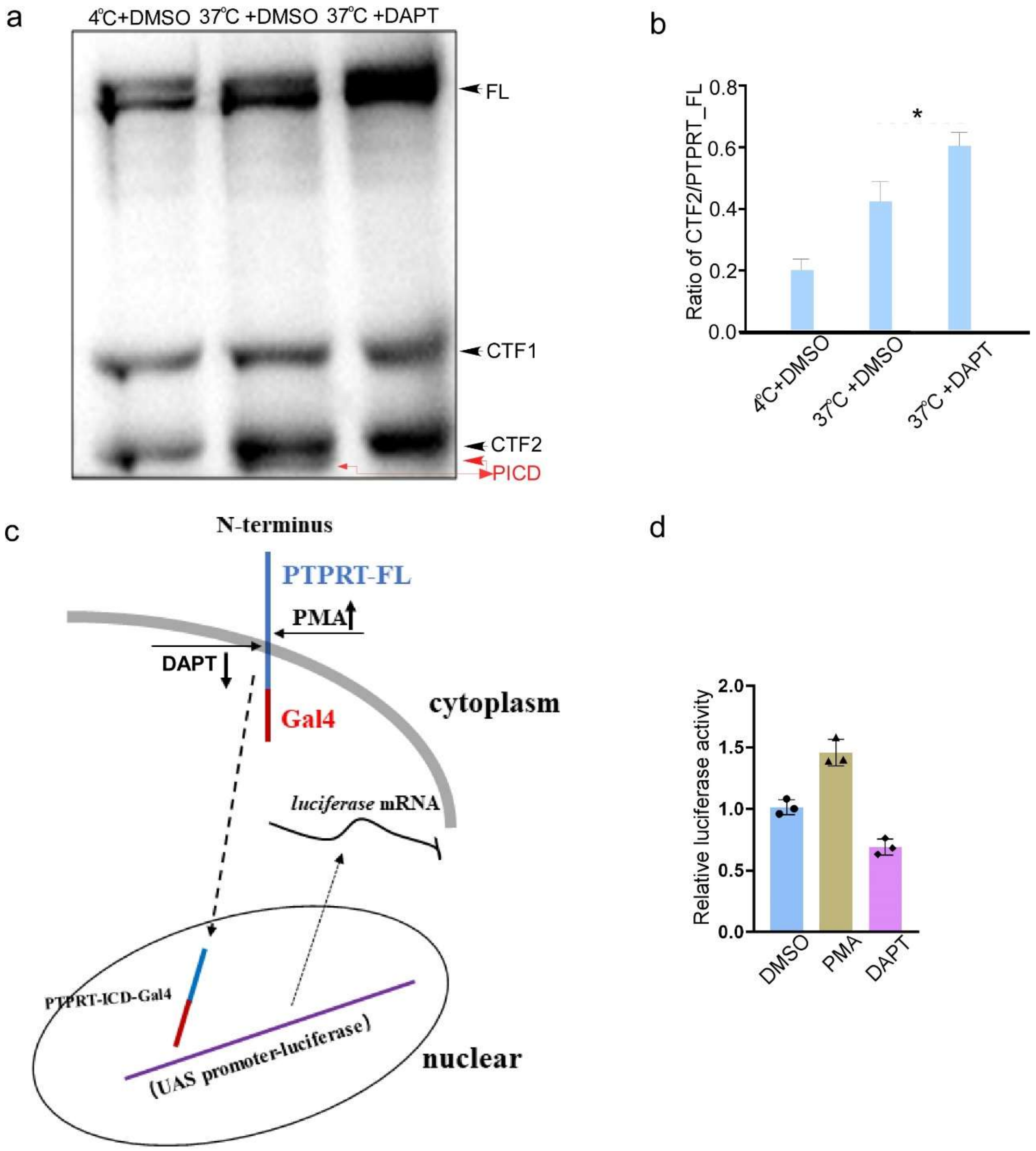


**Figure S1. Nuclear function of PTPRT intracellular domain induced by ADAM10 and presenilin 1/γ-secretase**

**a.** Protein extracts from HEK 293 cells with indicated pretreatments were assayed by Western blot for the presence of PTPRT, and its fragments CTF1 and CTF2 (PICD). **b.** The schematic of nuclear translocation and function of PTPRT intracellular domain induced by ADAM10 and presenilin 1/γ-secretase. **c.** The relative luciferase activities of luciferase reporters containing UAS promoter-luciferase, and HEK 293 cells transfected with PICD-Gal4. Results are presented as mean ±S.D. (n= 3 independent experiments, *p<0.01). **d.** Quantification of the immunobloted intensities of CTF2 shown in panel C). Error bars denote S.E.M. (n = 3 independent experiments; *, $P < 0.05$).

**Figure S2**

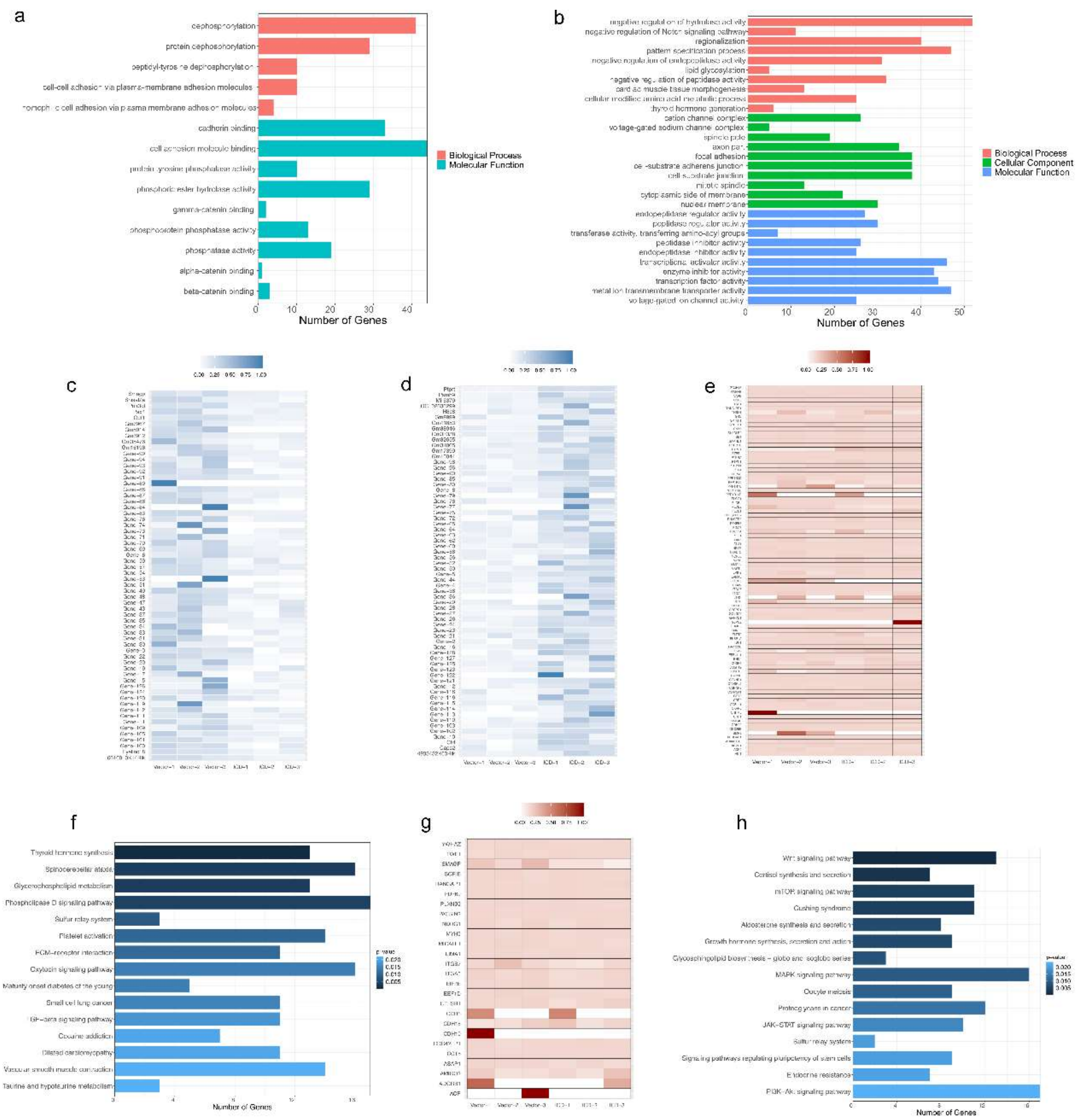

**Figure S2**

**Figure S2. PICD alter the expression of genes involved in neuronal circuit formation**

**a.** The GO annotation contains the *Ptprt* gene on chromosome 2. **b.** The number genes annotated to top 10 biological processes, cellular component, and molecular functions, respectively, on chromosome 2. **c-d.** Hierarchical clustering heat map of all down-regulated and up-regulated genes regulated by PICD. **e.** Heatmap of all gene expression has the same GO annotation with *ptprt* gene on chromosome 2 .**f.** The number of genes annotated to top 15 pathways on chromosome 2. **g.** Heatmap of expressions for genes have the same GO annotations with Smagp gene on chromosome 15. **h.** The number of genes annotated to top 15 pathways on chromosome 15.

**Figure S3**

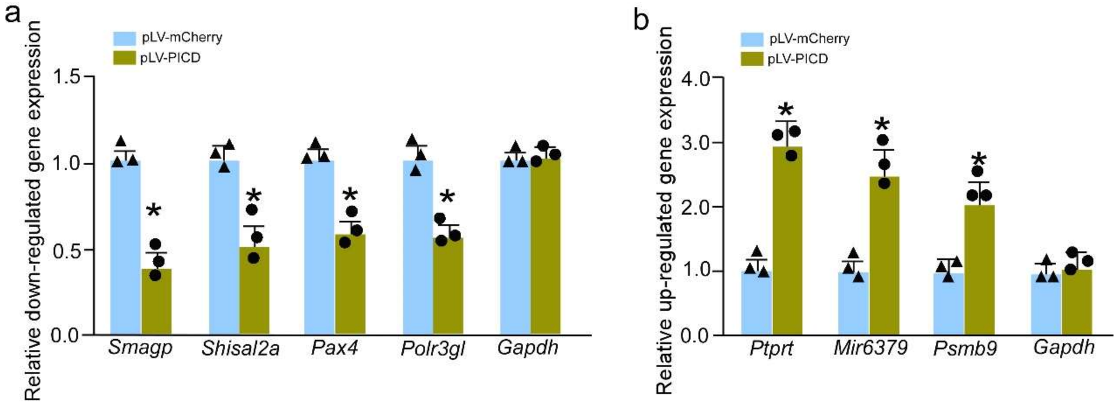


**Figure S3. Effect of PICD on targeted genes' mRNA expression**

**a.** Quantitative real-time PCR was performed with specific primers for downregulated mouse *Smagp*, *Shisal2a*, *Pax4* and *Polr3gl* mRNA expression. Total RNA was extracted from N2a cells transfected with pLV-mCherry or pLV-PICD-mCherry. Gapdh mRNA was used as internal control. Data represent mean ± S.E.M. of three independent experiments. * $P < 0.01$, unpaired *t*-test.**b.** Quantitative real-time PCR was performed with specific primers for upregulated mouse *Ptprt*, *Mir6379* and *Psmb9* mRNA expression. Total RNA was extracted from N2a cells transfected with pLV-mCherry or pLV-PICD-mCherry. *Gapdh* mRNA was used as internal control. Data represent mean ± S.E.M. of three independent experiments. * $P < 0.01$, unpaired *t*-test.

**Figure S4**

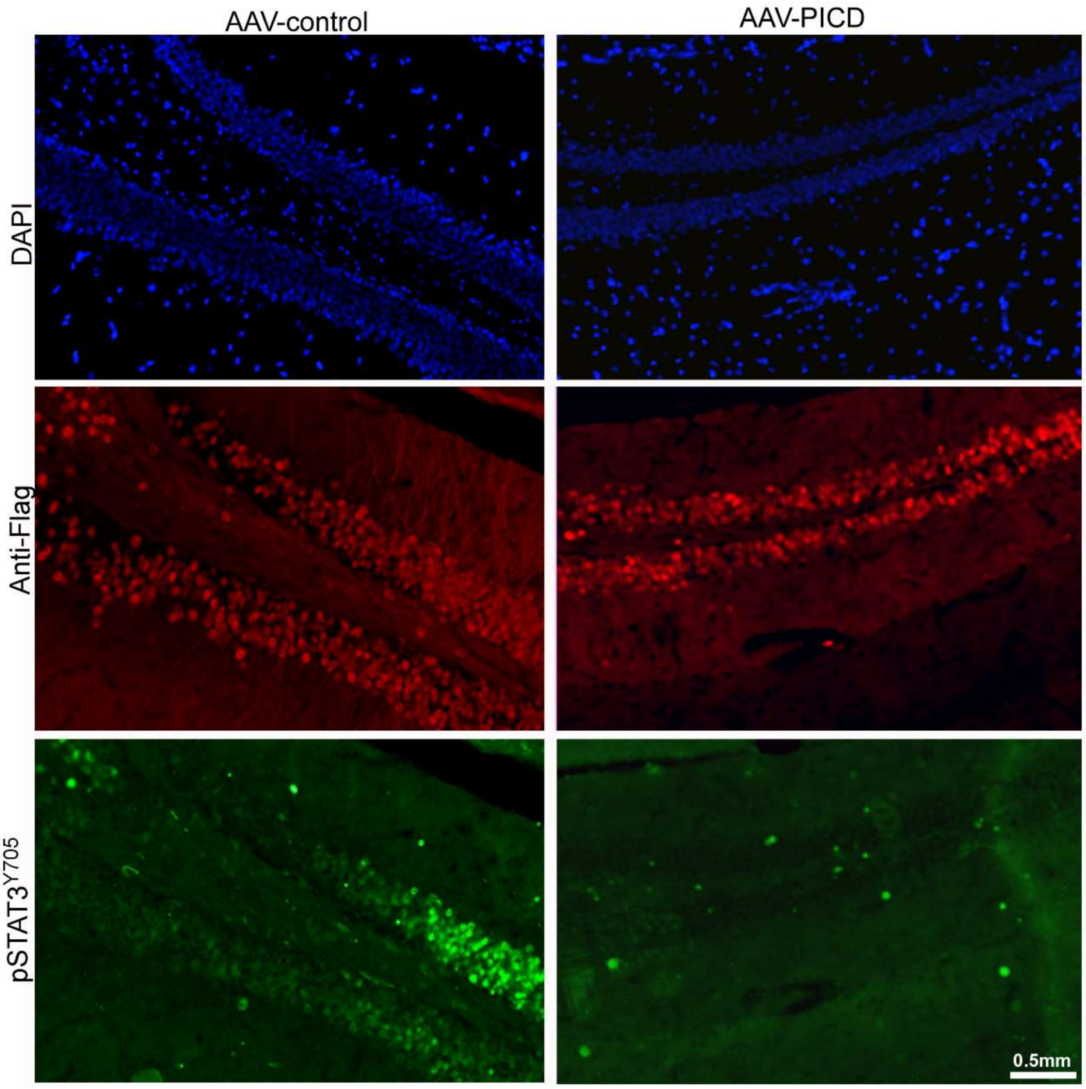


**Figure S4. Effect of AAV-PICD on pSTAT3$^{Y705}$ in APP/PS1 mice.** AAV-PICD could significantly decrease the pSTAT3$^{Y705}$ accumulation in APP/PS1 mouse brain.
